## Supplementary Information Fig S1 for "Structural basis for cell-type specific evolution of viral fitness by SARS-CoV-2"

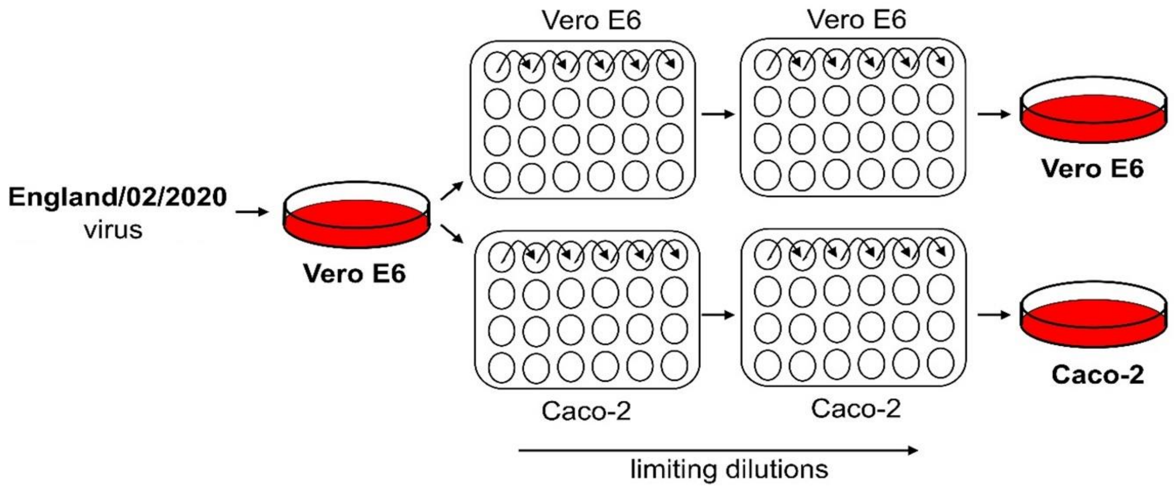


**Fig. S1 Isolation of the wildtype and BriSΔ viruses.** Schematic illustrating how the mixed population of SARS-CoV-2 viruses with wildtype (WT) and the "Bristol" variant BriSΔ were separated from each other by two rounds of limiting dilution in different cell types. After initial growth of the mixed virus population in Vero E6 cells, the virus titer was determined by qRT-PCR and serially diluted in 96 well plates until statistically just one virus particle was present in the wells with the highest dilution. In the wells with evidence of viral growth at the highest dilutions, the virus in the supernatant was assayed by site-specific PCR to determine if the virus was WT or the BriSΔ variant. This process was repeated to isolate a pure preparation of either virus which were then grown up to provide sufficient virus to sequence by dRNA-seq.
