## Supplementary Information Fig S2 for "Structural basis for cell-type specific evolution of viral fitness by SARS-CoV-2"

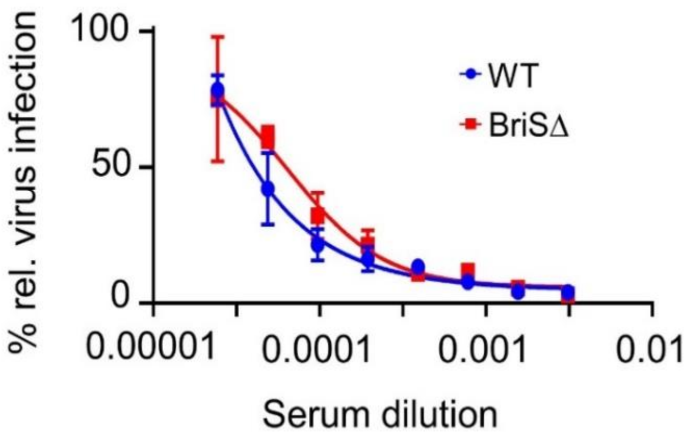


**Fig. S2 Human convalescent serum efficiently neutralizes WT and BriSΔ viruses.** Serum dilutions as indicated were utilized for virus neutralization using Vero E6 cells. WT: SARS-CoV-2 wildtype; BriSΔ : SARS-CoV-2 S deletion variant.
