## Supplementary Information Fig S3 for "Structural basis for cell-type specific evolution of viral fitness by SARS-CoV-2"

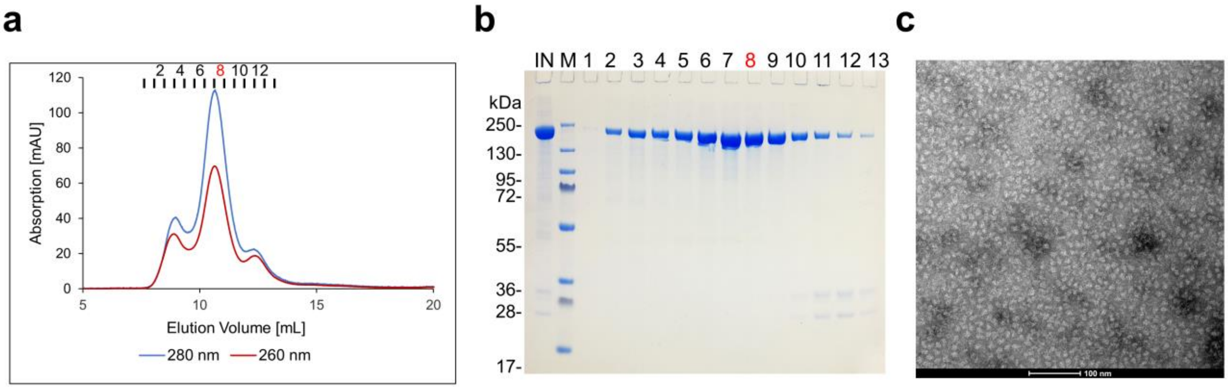


**Fig. S3 Purification and quality control of the BriSΔ glycoprotein. a**. Size-exclusion chromatography (SEC) of affinity-purified BriSΔ spike protein using a Superdex 200 column. Absorption was detected at 280 nm (blue line) and 260 nm (red line). Peak fractions are indicated. **b**. SDS PAGE analysis of the SEC fractions from panel A. lane 1: input fraction, lane 2: molecular weight marker, lanes 3-15: fractions 1 to 13 from SEC. **c**. Negative stain EM micrograph of SEC fraction 8 (scale bar: 100 nm).
