## Supplementary Information Fig S4 for "Structural basis for cell-type specific evolution of viral fitness by SARS-CoV-2"

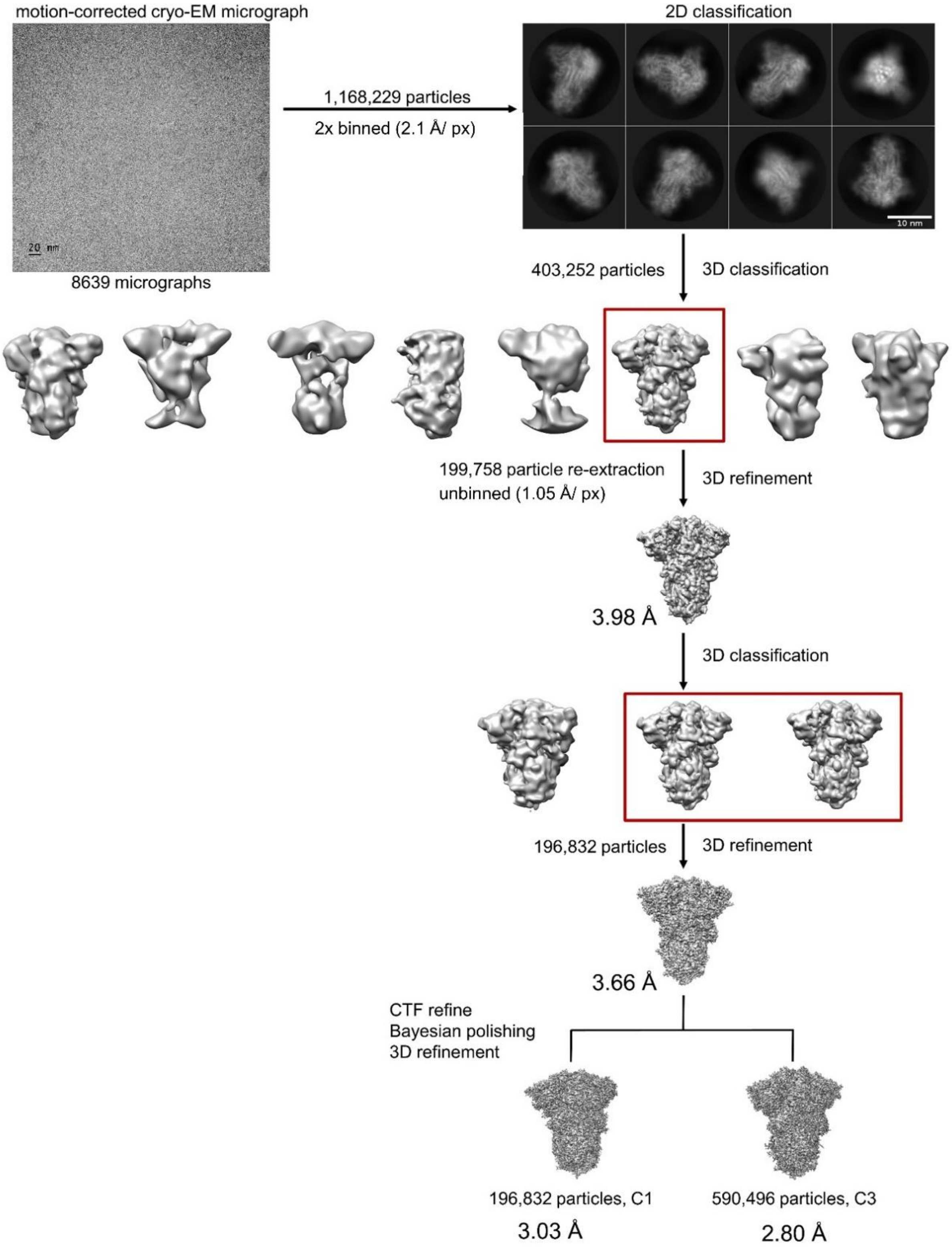


**Fig. S4 Cryo-EM image processing workflow**. A motion-corrected cryo-EM micrograph is shown (scale bar 20 nm, particles circled in red), reference-free 2D class averages (scale bar 10 nm), 3D classifications and refinements resulting in a not symmetrized C1 and a C3-symmetrized cryo-EM map.
