## Supplementary Information Fig S5 for "Structural basis for cell-type specific evolution of viral fitness by SARS-CoV-2"

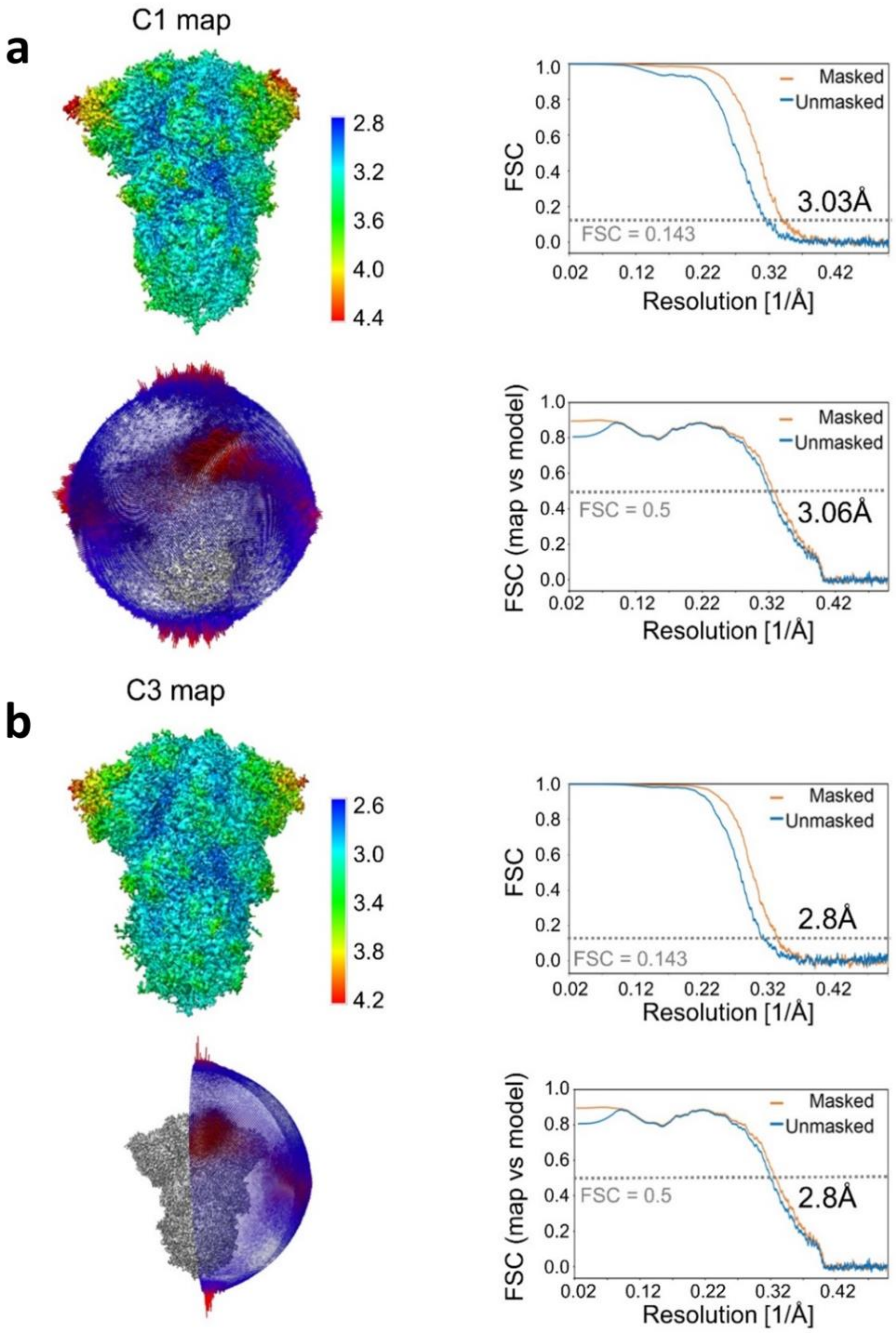


**Fig. S5 Cryo-EM structure validation.** Top left: Cryo-EM reconstruction colored according to the local resolution from a side view. Top right: Fourier Shell Correlation (FSC) curve after gold standard refinement. Below left: Orientation distribution of views that contributed to this map. Longer red rods represent orientations that comprise more particles. Below right: Cross-validation FSC curves for the refined model versus the final masked and unmasked maps, shown for **a**. the unsymmetrized C1 map and **b**. the C3-symmetrized map.
