## Supplementary Information Fig S6 for "Structural basis for cell-type specific evolution of viral fitness by SARS-CoV-2"

a

b

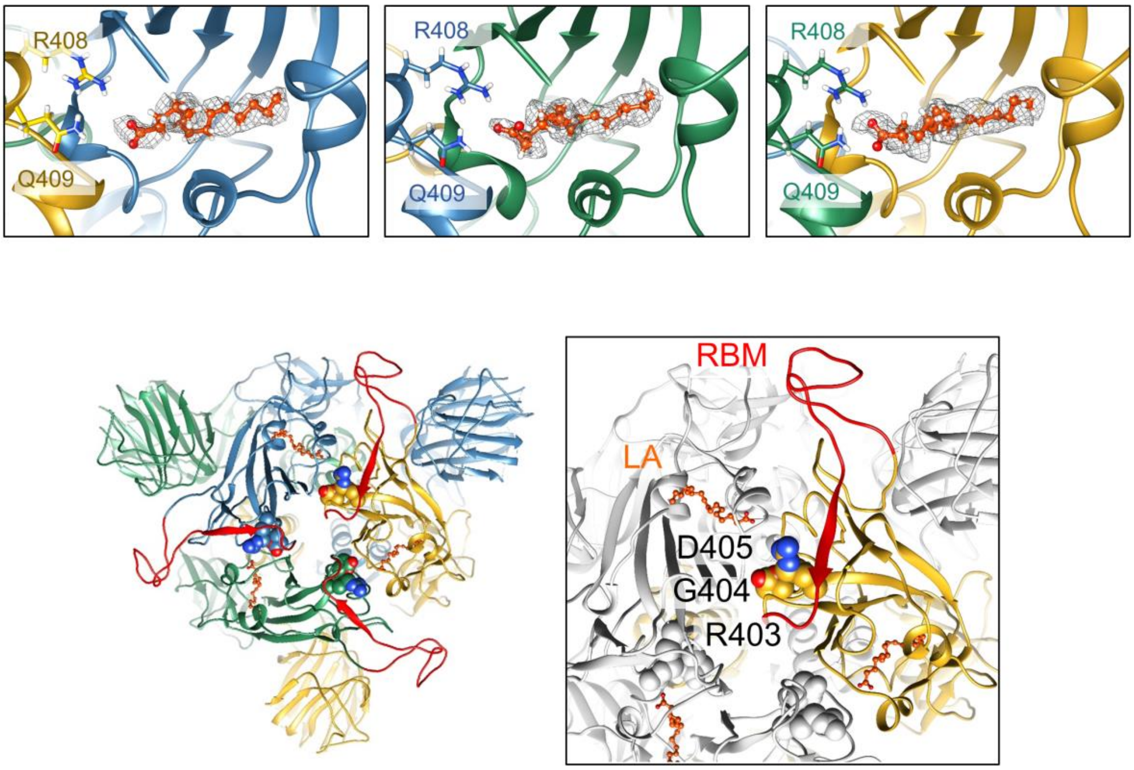


**Fig. S6 BriSΔ RBD structural organization. a**. LA binding in each bipartite free fatty acid binding pocket in the unsymmetrized C1 structure is shown. EM density is shown as grey mesh. **b**. The positions of LA, the receptor-binding motif (RBM) and the arginine-glycine-aspartate tripeptide (RGD) motif are shown from a top view on the left. LA is colored in orange, RBM is colored in red. RGD is shown as spheres. A section is depicted in a zoom-in on the right. LA, RBM and RGD are indicated.
