## Supplementary Information Fig S7 for "Structural basis for cell-type specific evolution of viral fitness by SARS-CoV-2"

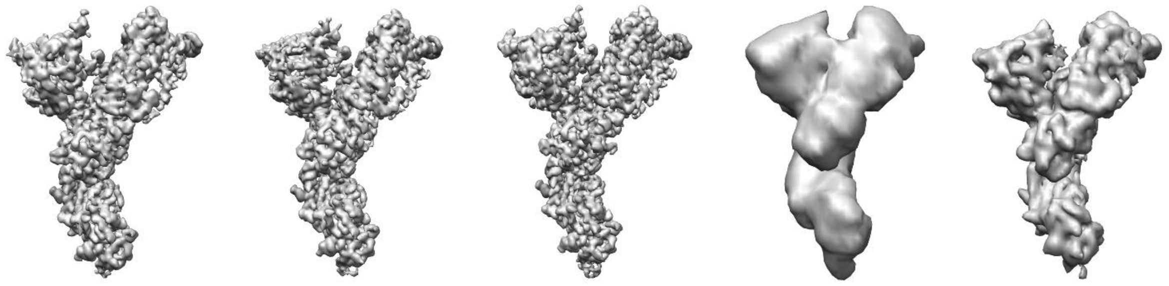

| Class 1 | Class 2 | Class 3 | Class 4 | Class 5 |
|---|---|---|---|---|
| 111,217 particles | 214,977 particles | 235,538 particles | 130 particles | 28,634 particles |
| 18.8 % | 36.4 % | 39.9 % | 0.02 % | 4.8 % |

**Fig. S7 Masked 3D classification.** Masked 3D classification focusing of individual chains within the S trimer into 5 classes. Class 1-3 comprised 95% of all particles. These classes all present LA-bound RBDs. The other 3D classes did not reach sufficient resolution to determine the presence or absence of LA in the RBD reconstructions.
