## Supplementary Information Fig S8 for "Structural basis for cell-type specific evolution of viral fitness by SARS-CoV-2"

**a**

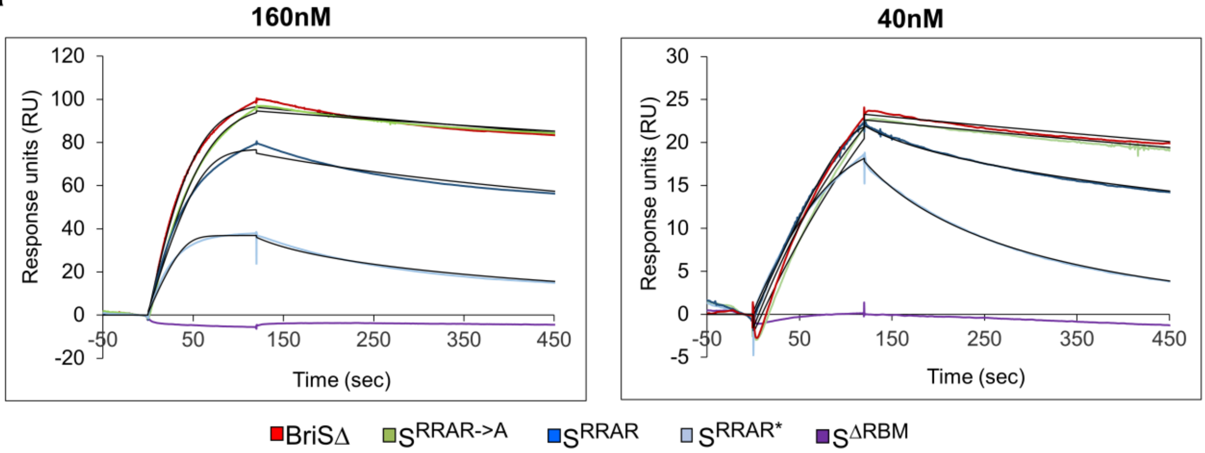


**b**

| | $K_D$ (nM) | $k_{on}$ $M^{-1}$ $s^{-1}$ | $k_{off}$ = $s^{-1}$ |
|---|---|---|---|
| BriSΔ | 1.8 ± 0.34 | 2.22 ± 0.49 x $10^5$ | 3.9 ± 0.4 x $10^{-4}$ |
| $S^{RRAR\text{->}A}$ | 1.4 ± 0.3 | 2.5 ± 1.2 x $10^5$ | 3.6 ± 2.2 x $10^{-4}$ |
| $S^{RRAR}$ | 2.51 ± 0.06 | 4.55 ± 0.04 x $10^5$ | 11.4 ± 0.28 x $10^{-4}$ |
| $S^{RRAR*}$ | 12.2 ± 4.36 | 6.33 ± 2.18 x $10^5$ | 70.69 ± 5.94 x $10^{-4}$ |

**Fig. S8 Spike proteins binding to ACE2 by Surface Plasmon Resonance (SPR). a**. Spike proteins BriSΔ, $S^{RRAR\text{->}A}$, $S^{RRAR}$ and furin-cleaved $S^{RRAR*}$ were analyzed for binding to ACE2 receptor immobilized on a streptavidin-coated sensor chip. Sensorgrams for representative concentrations (160 nM and 40 nM) are shown including $S^{\Delta RBM}$ lacking the receptor-binding motif as control. $K_D$ values, $k_{on}$ and $k_{off}$ are listed in **b**. Values for $S^{RRAR\text{->}A}$ are from {Toelzer, 2020 #90}.
