## Supplementary Information Fig S9 for "Structural basis for cell-type specific evolution of viral fitness by SARS-CoV-2"

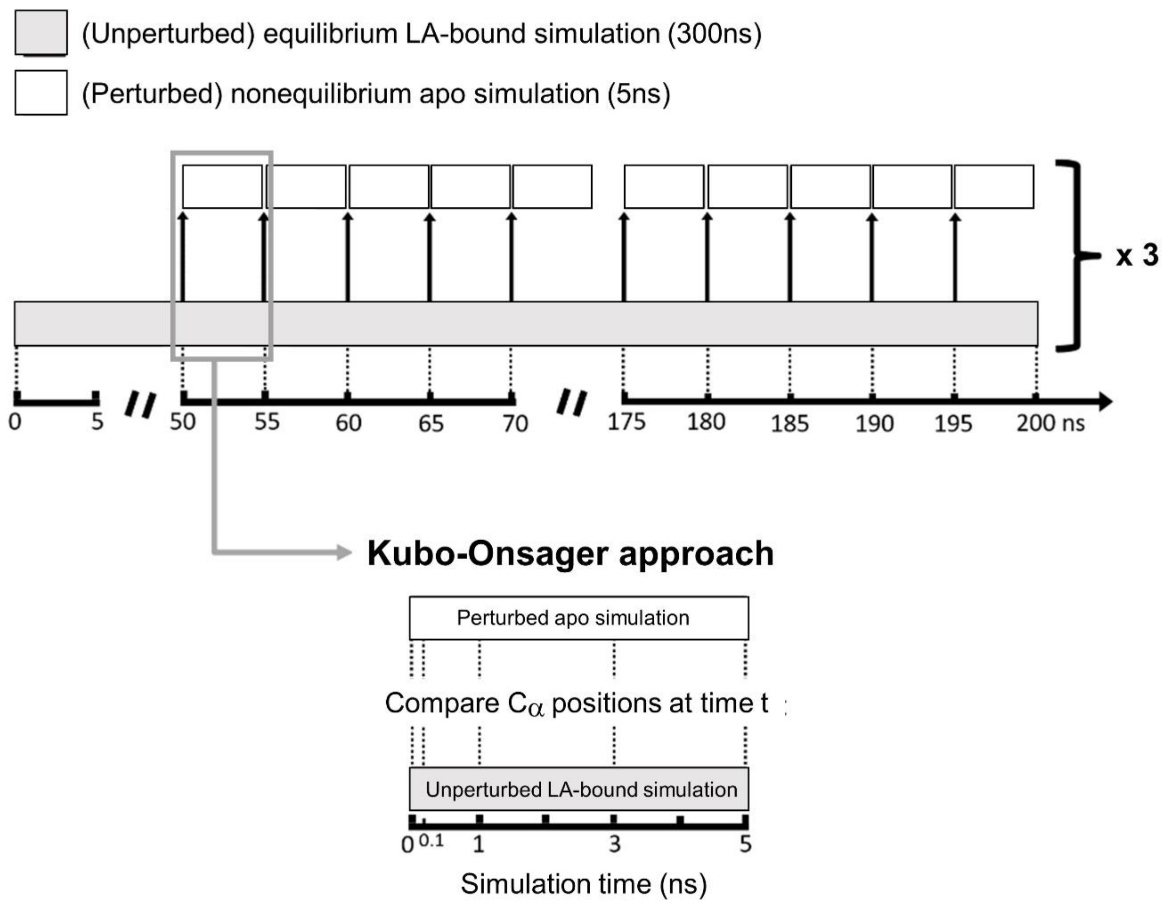


**Fig. S9 Nonequilibrium simulations of wildtype S and BriSΔ proteins.** A schematic description of the procedure used to set up and analyze the nonequilibrium MD simulations. From the cryo-EM structure of the S proteins with LA bound, three equilibrium MD simulations, 200 ns each, were performed. These equilibrium simulations (indicated by rectangles filled in grey) were used to generate starting conformations for the nonequilibrium simulations (rectangles filled in white). Conformations were extracted every five nanoseconds from the equilibrated part of each LA-bound simulation (from 50-300 ns), and LA was removed. Each nonequilibrium simulation was run for five nanoseconds. The Kubo-Onsager approach was used to extract the response of the system to LA removal (bottom panel). For each pair of unperturbed LA-bound and perturbed apo simulations, the positional deviations of each Cα at equivalent times (namely 0, 0.1, 1, 3 and 5 ns) were determined and averaged over all 90 simulations.
