## Supplementary Information Fig S10 for "Structural basis for cell-type specific evolution of viral fitness by SARS-CoV-2"

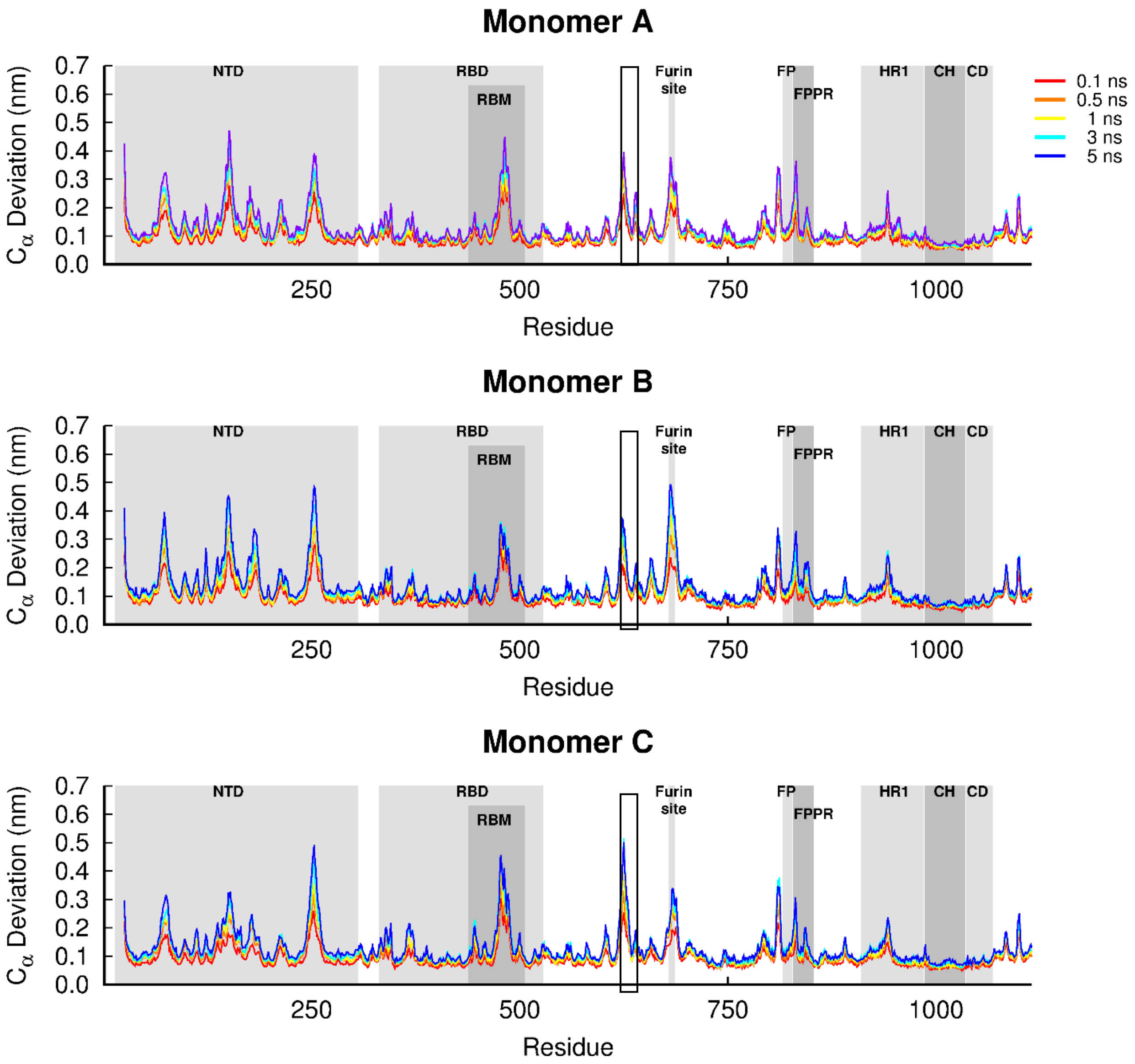


**Fig. S10 Average Cα-positional deviation in the five nanoseconds after removing LA from wildtype S protein**. The average deviations were calculated using the Kubo-Onsager approach for the pairwise comparison between the nonequilibrium apo and equilibrium LA-bound simulations. The positions of important structural motifs are highlighted in grey, namely N-terminal domain (NTD), receptor-binding domain (RBD), receptor-binding motif (RBM), fusion peptide (FP), fusion peptide proximal region (FPPR), heptad repeat 1 (HR1), central helix (CH) and connector domain (CD). The V622-L629 region adjacent to R634 is boxed in black.
