## Supplementary Information Fig S11 for "Structural basis for cell-type specific evolution of viral fitness by SARS-CoV-2"

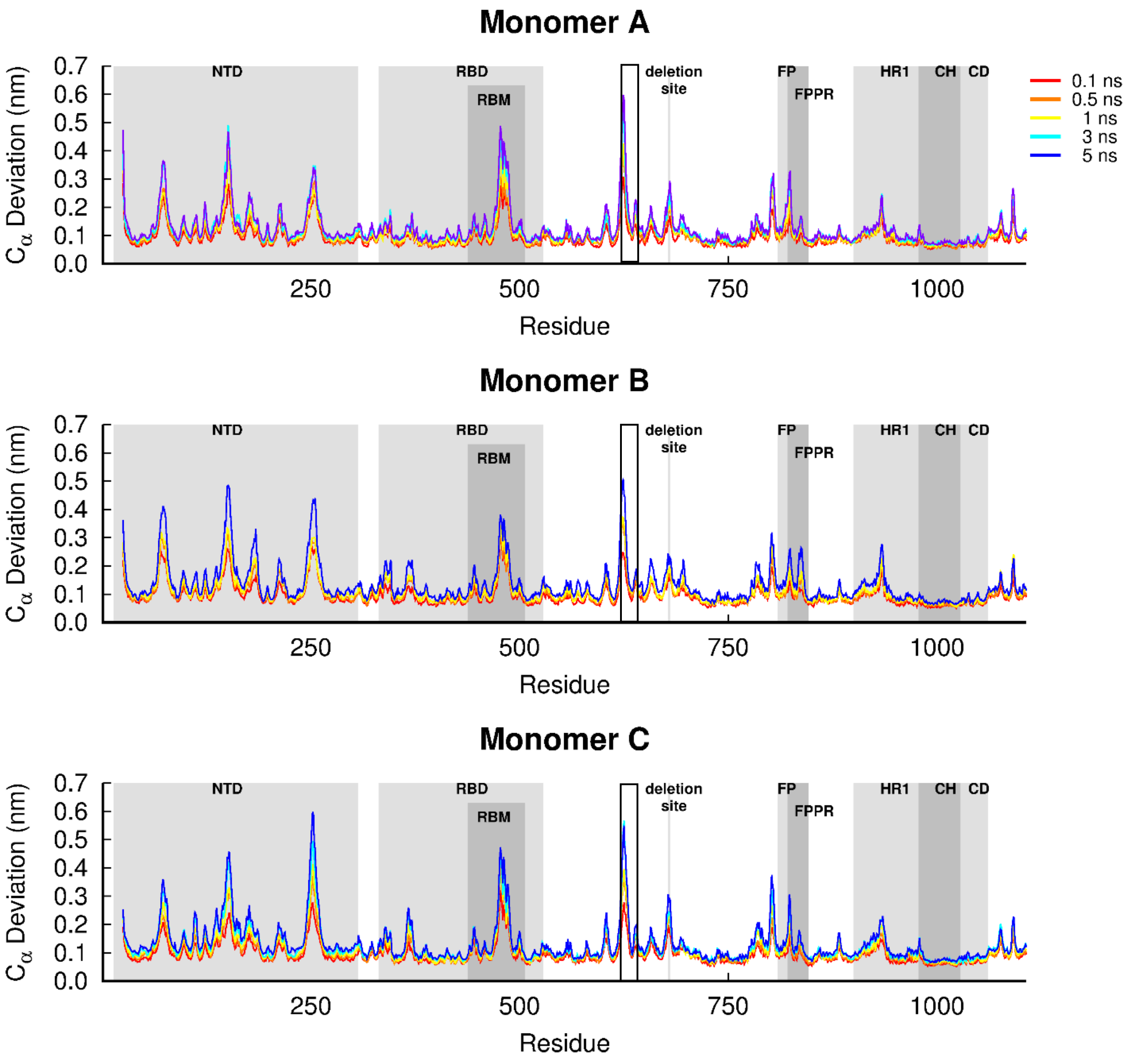


**Fig. S11 Average Cα-positional deviation in the five nanoseconds after removing LA from BriSΔ protein.** The average deviations were calculated using the Kubo-Onsager approach for the pairwise comparison between the nonequilibrium apo and equilibrium LA-bound simulations. The positions of relevant structural motifs are highlighted in grey, namely the N-terminal domain (NTD), receptor-binding domain (RBD), receptor-binding motif (RBM), fusion peptide (FP), fusion peptide proximal region (FPPR), heptad repeat 1 (HR1), central helix (CH), connector domain (CD). The V622-L629 region adjacent to R634 is boxed in black.
