## Supplementary Information Table S1 for "Structural basis for cell-type specific evolution of viral fitness by SARS-CoV-2"

**Table S1. Cryo-EM data collection and refinement statistics.**

| | C1 | C3 |
|---|---|---|
| **Data collection and processing** | | |
| Voltage (kV) | 200 | 200 |
| Magnification (nominal) | 130,000 | 130,000 |
| Pixel size (Å / px) | 1.05 (0.525) | 1.05 (0.525) |
| Flux ($e^-$ /pix/sec) | 6.4 | 6.4 |
| Frames per exposure | 55 | 55 |
| Exposure ($e^-$ / Å$^2$) | 1.16 | 1.16 |
| Defocus range (µm) | -0.8 to -2.0 | -0.8 to -2.0 |
| Micrographs collected | 9,519 | 9,519 |
| Particles final | 196,832 | 590,496 |
| Map sharpening B-factor (Å$^2$) | -97.6 | -106.7 |
| Masked resolution at 0.143 FSC (Å) | 3.03 | 2.8 |
| **Refinement** | | |
| Composition | | |
| Amino acids | 3066 | 3036 |
| Glycans | 30 | 33 |
| Ligands | 3 | 3 |
| RMSD bonds (Å) | 0.003 | 0.004 |
| RMSD angles (º) | 0.564 | 0.555 |
| Mean B-factors (Å$^2$) | | |
| Amino acids | 18.75 | 29.26 |
| Ligands | 33.61 | 41.97 |
| Ramachandran | | |
| Favored (%) | 94.06 | 95.22 |
| Allowed (%) | 5.84 | 4.78 |
| Outliers (%) | 0.1 | 0.00 |
| Rotamer outliers (%) | 0.64 | 0.23 |
| Clash score | 2.77 | 2.49 |
| C-beta outliers (%) | 0.00 | 0.00 |
| CaBLAM outliers (%) | 3.14 | 2.51 |
| CC (mask) | 0.82 | 0.80 |
| MolProbity score | 1.46 | 1.36 |
| EMRinger score | 3.77 | 3.95 |
| Model resolution (Å), 0.5 FSC threshold | 3.0 | 2.8 |
