## Supplementary Information Table S2 for "Structural basis for cell-type specific evolution of viral fitness by SARS-CoV-2"

**Table S2. N-linked glycosylation sites in recombinant SARS-CoV-2 S proteins.**

| **WT SARS-CoV-2 S *** | **Recombinant SARS-CoV-2 S expressed in** | | |
|---|---|---|---|
| **Predicted (UNIPROT)** | **Freestyle 293F** [8] | **Hi5** [23] | **Hi5** (BrisΔ, this study) |
| $N_{17}$LT | | $N_{17}$LT | |
| $N_{61}$VT | $N_{61}$VT | $N_{61}$VT | $N_{61}$VT |
| $N_{74}$GT | | | |
| $N_{121}$NA # | | | |
| $N_{122}$AT | $N_{122}$AT | $N_{122}$AT | $N_{122}$AT |
| $N_{149}$KS | | | |
| $N_{165}$CT | $N_{165}$CT | $N_{165}$CT | $N_{165}$CT |
| $N_{234}$IT | $N_{234}$IT | $N_{234}$IT | $N_{234}$IT |
| $N_{282}$GT | $N_{282}$GT | $N_{282}$GT | $N_{282}$GT |
| $N_{331}$IT | $N_{331}$IT | $N_{331}$IT | $N_{331}$IT |
| $N_{343}$AT | $N_{343}$AT | $N_{343}$AT | $N_{343}$AT |
| $N_{370}$SA # | | | |
| $N_{603}$TS | $N_{603}$TS | | |
| $N_{616}$CT | $N_{616}$CT | $N_{616}$CT | $N_{616}$CT |
| $N_{657}$NS | $N_{657}$NS | | $N_{657}$NS |
| $N_{709}$NS | $N_{709}$NS | $N_{706}$NS | $N_{701}$NS |
| $N_{717}$FT | $N_{717}$FT | $N_{714}$FT | $N_{709}$FT |
| $N_{801}$FS | $N_{801}$FS | $N_{798}$FS | $N_{793}$FS |
| $N_{1074}$FT | $N_{1074}$FT | $N_{1071}$FT | $N_{1066}$FT |
| $N_{1098}$GT | $N_{1098}$GT | $N_{1095}$GT | $N_{1090}$GT |
| $N_{1134}$NT | $N_{1134}$NT | $N_{1131}$NT | $N_{1126}$NT |
| $N_{1158}$HT | | | |
| $N_{1173}$AS | | | |
| $N_{1194}$ES | | | |

*Sites lacking glycosylation in cryo-EM maps are omitted (boxes colored in grey).*

***** *YP_009724390.1*

### *Predicted based on SARS-CoV-2 S protein*[8]
