## Supplementary Information Table S2 for "Structural basis for cell-type specific evolution of viral fitness by SARS-CoV-2"

**Table S3. Alignment of SARS-CoV-2 S protein sequences used in this study**

```
             1        10        20        30        40        50        60
S WT         MFVFLVLLPLVSSQCVNLTTRTQLPPAYTNSFTRGVYYPDKVFRSSVLHSTQDLFLPFFS
BriSΔ        MFVFLVLLPLVSSQCVNLTTRTQLPPAYTNSFTRGVYYPDKVFRSSVLHSTQDLFLPFFS
S RRAR->A    MFVFLVLLPLVSSQCVNLTTRTQLPPAYTNSFTRGVYYPDKVFRSSVLHSTQDLFLPFFS
S RRAR       MFVFLVLLPLVSSQCVNLTTRTQLPPAYTNSFTRGVYYPDKVFRSSVLHSTQDLFLPFFS
S RRAR*      MFVFLVLLPLVSSQCVNLTTRTQLPPAYTNSFTRGVYYPDKVFRSSVLHSTQDLFLPFFS
S ΔRMB       MFVFLVLLPLVSSQCVNLTTRTQLPPAYTNSFTRGVYYPDKVFRSSVLHSTQDLFLPFFS

                      70        80        90       100       110       120
S WT         NVTWFHAIHVSGTNGTKRFDNPVLPFNDGVYFASTEKSNIIRGWIFGTTLDSKTQSLLIV
BriSΔ        NVTWFHAIHVSGTNGTKRFDNPVLPFNDGVYFASTEKSNIIRGWIFGTTLDSKTQSLLIV
S RRAR->A    NVTWFHAIHVSGTNGTKRFDNPVLPFNDGVYFASTEKSNIIRGWIFGTTLDSKTQSLLIV
S RRAR       NVTWFHAIHVSGTNGTKRFDNPVLPFNDGVYFASTEKSNIIRGWIFGTTLDSKTQSLLIV
S RRAR*      NVTWFHAIHVSGTNGTKRFDNPVLPFNDGVYFASTEKSNIIRGWIFGTTLDSKTQSLLIV
S ΔRMB       NVTWFHAIHVSGTNGTKRFDNPVLPFNDGVYFASTEKSNIIRGWIFGTTLDSKTQSLLIV

                     130       140       150       160       170       180
S WT         NNATNVVIKVCEFQFCNDPFLGVYYHKNNKSWMESEFRVYSSANNCTFEYVSQPFLMDLE
BriSΔ        NNATNVVIKVCEFQFCNDPFLGVYYHKNNKSWMESEFRVYSSANNCTFEYVSQPFLMDLE
S RRAR->A    NNATNVVIKVCEFQFCNDPFLGVYYHKNNKSWMESEFRVYSSANNCTFEYVSQPFLMDLE
S RRAR       NNATNVVIKVCEFQFCNDPFLGVYYHKNNKSWMESEFRVYSSANNCTFEYVSQPFLMDLE
S RRAR*      NNATNVVIKVCEFQFCNDPFLGVYYHKNNKSWMESEFRVYSSANNCTFEYVSQPFLMDLE
S ΔRMB       NNATNVVIKVCEFQFCNDPFLGVYYHKNNKSWMESEFRVYSSANNCTFEYVSQPFLMDLE

                     190       200       210       220       230       240
S WT         GKQGNFKNLREFVFKNIDGYFKIYSKHTPINLVRDLPQGFSALEPLVDLPIGINITRFQT
BriSΔ        GKQGNFKNLREFVFKNIDGYFKIYSKHTPINLVRDLPQGFSALEPLVDLPIGINITRFQT
S RRAR->A    GKQGNFKNLREFVFKNIDGYFKIYSKHTPINLVRDLPQGFSALEPLVDLPIGINITRFQT
S RRAR       GKQGNFKNLREFVFKNIDGYFKIYSKHTPINLVRDLPQGFSALEPLVDLPIGINITRFQT
S RRAR*      GKQGNFKNLREFVFKNIDGYFKIYSKHTPINLVRDLPQGFSALEPLVDLPIGINITRFQT
S ΔRMB       GKQGNFKNLREFVFKNIDGYFKIYSKHTPINLVRDLPQGFSALEPLVDLPIGINITRFQT

                     250       260       270       280       290       300
S WT         LLALHRSYLTPGDSSSGWTAGAAAYYVGYLQPRTFLLKYNENGTITDAVDCALDPLSETK
BriSΔ        LLALHRSYLTPGDSSSGWTAGAAAYYVGYLQPRTFLLKYNENGTITDAVDCALDPLSETK
S RRAR->A    LLALHRSYLTPGDSSSGWTAGAAAYYVGYLQPRTFLLKYNENGTITDAVDCALDPLSETK
S RRAR       LLALHRSYLTPGDSSSGWTAGAAAYYVGYLQPRTFLLKYNENGTITDAVDCALDPLSETK
S RRAR*      LLALHRSYLTPGDSSSGWTAGAAAYYVGYLQPRTFLLKYNENGTITDAVDCALDPLSETK
S ΔRMB       LLALHRSYLTPGDSSSGWTAGAAAYYVGYLQPRTFLLKYNENGTITDAVDCALDPLSETK

                     310       320       330       340       350       360
S WT         CTLKSFTVEKGIYQTSNFRVQPTESIVRFPNITNLCPFGEVFNATRFASVYAWNRKRISN
BriSΔ        CTLKSFTVEKGIYQTSNFRVQPTESIVRFPNITNLCPFGEVFNATRFASVYAWNRKRISN
S RRAR->A    CTLKSFTVEKGIYQTSNFRVQPTESIVRFPNITNLCPFGEVFNATRFASVYAWNRKRISN
S RRAR       CTLKSFTVEKGIYQTSNFRVQPTESIVRFPNITNLCPFGEVFNATRFASVYAWNRKRISN
S RRAR*      CTLKSFTVEKGIYQTSNFRVQPTESIVRFPNITNLCPFGEVFNATRFASVYAWNRKRISN
S ΔRMB       CTLKSFTVEKGIYQTSNFRVQPTESIVRFPNITNLCPFGEVFNATRFASVYAWNRKRISN

                     370       380       390       400       410       420
S WT         CVADYSVLYNSASFSTFKCYGVSPTKLNDLCFTNVYADSFVIRGDEVRQIAPGQTGKIAD
BriSΔ        CVADYSVLYNSASFSTFKCYGVSPTKLNDLCFTNVYADSFVIRGDEVRQIAPGQTGKIAD
S RRAR->A    CVADYSVLYNSASFSTFKCYGVSPTKLNDLCFTNVYADSFVIRGDEVRQIAPGQTGKIAD
S RRAR       CVADYSVLYNSASFSTFKCYGVSPTKLNDLCFTNVYADSFVIRGDEVRQIAPGQTGKIAD
S RRAR*      CVADYSVLYNSASFSTFKCYGVSPTKLNDLCFTNVYADSFVIRGDEVRQIAPGQTGKIAD
S ΔRMB       CVADYSVLYNSASFSTFKCYGVSPTKLNDLCFTNVYADSFVIRGDEVRQIAPGQTGKIAD

                     430       440       450       460       470       480
S WT         YNYKLPDDFTGCVIAWNSNNLDSKVGGNYNYLYRLFRKSNLKPFERDISTEIYQAGSTPC
BriSΔ        YNYKLPDDFTGCVIAWNSNNLDSKVGGNYNYLYRLFRKSNLKPFERDISTEIYQAGSTPC
S RRAR->A    YNYKLPDDFTGCVIAWNSNNLDSKVGGNYNYLYRLFRKSNLKPFERDISTEIYQAGSTPC
S RRAR       YNYKLPDDFTGCVIAWNSNNLDSKVGGNYNYLYRLFRKSNLKPFERDISTEIYQAGSTPC
S RRAR*      YNYKLPDDFTGCVIAWNSNNLDSKVGGNYNYLYRLFRKSNLKPFERDISTEIYQAGSTPC
S ΔRMB       YNYKLPDDFTGCVIAWNSNNLDSS.GGSYNYLARG.........................
```

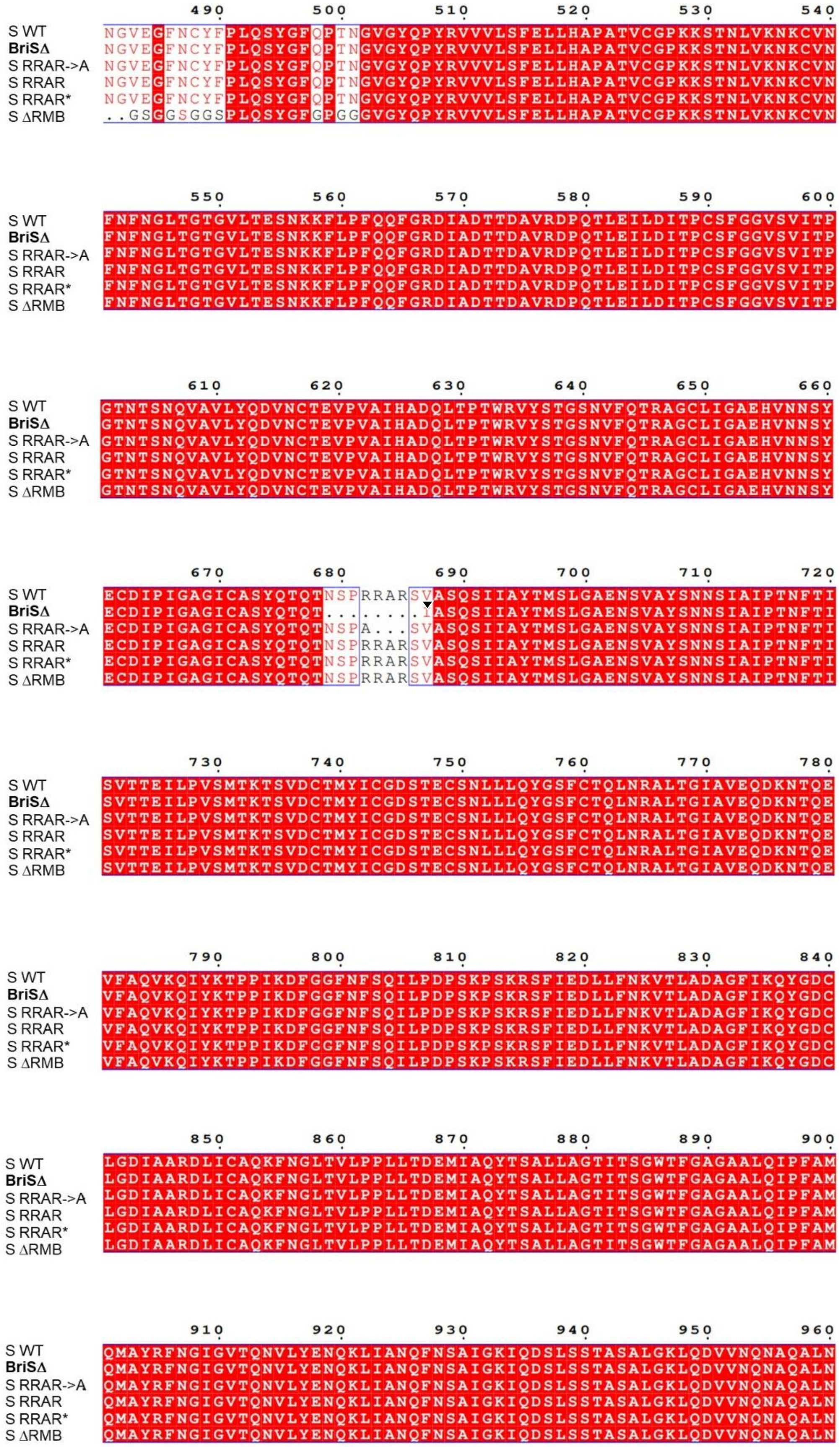


* *Furin cleavage in* $S^{RRAR*}$ *is marked with a triangle filled in black*

```
              970       980       990       1000      1010      1020
S WT          TLVKQLSSNFGAISSVLNDILSRLDKVEAEVQIDRLITGRLQSLQTYVTQQLIRAAEIRA
BriSΔ         TLVKQLSSNFGAISSVLNDILSRLDKVEAEVQIDRLITGRLQSLQTYVTQQLIRAAEIRA
S RRAR->A     TLVKQLSSNFGAISSVLNDILSRLDKVEAEVQIDRLITGRLQSLQTYVTQQLIRAAEIRA
S RRAR        TLVKQLSSNFGAISSVLNDILSRLDPPEAEVQIDRLITGRLQSLQTYVTQQLIRAAEIRA
S RRAR*       TLVKQLSSNFGAISSVLNDILSRLDPPEAEVQIDRLITGRLQSLQTYVTQQLIRAAEIRA
S ΔRMB        TLVKQLSSNFGAISSVLNDILSRLDPPEAEVQIDRLITGRLQSLQTYVTQQLIRAAEIRA
```

```
              1030      1040      1050      1060      1070      1080
S WT          SANLAATKMSECVLGQSKRVDFCGKGYHLMSFPQSAPHGVVFLHVTYVPAQEKNFTTAPA
BriSΔ         SANLAATKMSECVLGQSKRVDFCGKGYHLMSFPQSAPHGVVFLHVTYVPAQEKNFTTAPA
S RRAR->A     SANLAATKMSECVLGQSKRVDFCGKGYHLMSFPQSAPHGVVFLHVTYVPAQEKNFTTAPA
S RRAR        SANLAATKMSECVLGQSKRVDFCGKGYHLMSFPQSAPHGVVFLHVTYVPAQEKNFTTAPA
S RRAR*       SANLAATKMSECVLGQSKRVDFCGKGYHLMSFPQSAPHGVVFLHVTYVPAQEKNFTTAPA
S ΔRMB        SANLAATKMSECVLGQSKRVDFCGKGYHLMSFPQSAPHGVVFLHVTYVPAQEKNFTTAPA
```

```
              1090      1100      1110      1120      1130      1140
S WT          ICHDGKAHFPREGVFVSNGTHWFVTQRNFYEPQIITTDNTFVSGNCDVVIGIVNNTVYDP
BriSΔ         ICHDGKAHFPREGVFVSNGTHWFVTQRNFYEPQIITTDNTFVSGNCDVVIGIVNNTVYDP
S RRAR->A     ICHDGKAHFPREGVFVSNGTHWFVTQRNFYEPQIITTDNTFVSGNCDVVIGIVNNTVYDP
S RRAR        ICHDGKAHFPREGVFVSNGTHWFVTQRNFYEPQIITTDNTFVSGNCDVVIGIVNNTVYDP
S RRAR*       ICHDGKAHFPREGVFVSNGTHWFVTQRNFYEPQIITTDNTFVSGNCDVVIGIVNNTVYDP
S ΔRMB        ICHDGKAHFPREGVFVSNGTHWFVTQRNFYEPQIITTDNTFVSGNCDVVIGIVNNTVYDP
```

```
              1150      1160      1170      1180      1190      1200
S WT          LQPELDSFKEELDKYFKNHTSPDVDLGDISGINASVVNIQKEIDRLNEVAKNLNESLIDL
BriSΔ         LQPELDSFKEELDKYFKNHTSPDVDLGDISGINASVVNIQKEIDRLNEVAKNLNESLIDL
S RRAR->A     LQPELDSFKEELDKYFKNHTSPDVDLGDISGINASVVNIQKEIDRLNEVAKNLNESLIDL
S RRAR        LQPELDSFKEELDKYFKNHTSPDVDLGDISGINASVVNIQKEIDRLNEVAKNLNESLIDL
S RRAR*       LQPELDSFKEELDKYFKNHTSPDVDLGDISGINASVVNIQKEIDRLNEVAKNLNESLIDL
S ΔRMB        LQPELDSFKEELDKYFKNHTSPDVDLGDISGINASVVNIQKEIDRLNEVAKNLNESLIDL
```

```
              1210      1220      1230                1240
S WT          QELGKYEQYIKWPWYIWLGFIA...GLIAIVMV...........TIMLCCMTSCCSCLKG
BriSΔ         QELGKYEQG.....GGGSGGGGSGSGYIPEAPRDGQAYVRKDGEWVLLSTFLGGGGG.SG
S RRAR->A     QELGKYEQYIKWPSGRLVPRGSPGSGYIPEAPRDGQAYVRKDGEWVLLSTFLGHHHH.HH
S RRAR        QELGKYEQG.....GGGSGGGGSGSGYIPEAPRDGQAYVRKDGEWVLLSTFLGGGGG.SG
S RRAR*       QELGKYEQG.....GGGSGGGGSGSGYIPEAPRDGQAYVRKDGEWVLLSTFLGGGGG.SG
S ΔRMB        QELGKYEQG.....GGGSGGGGSGSGYIPEAPRDGQAYVRKDGEWVLLSTFLGGGGG.SG
```

```
              1250      1260      1270
S WT          CCSCGSCCKFDEDDSEPVLKGVKLHYT...
BriSΔ         GGGSGSEQKLISEEDLGSGSGSHHHHHHHH
S RRAR->A     ..............................
S RRAR        GGGSGSEQKLISEEDLGSGSGSHHHHHHHH
S RRAR*       GGGSGSEQKLISEEDLGSGSGSHHHHHHHH
S ΔRMB        GGGSGSEQKLISEEDLGSGSGSHHHHHHHH
```
